# Integrating geospatial tools in mapping forest fire severity and burned areas in the Western Usambara Mountain Forests, Lushoto, Tanzania

**DOI:** 10.1101/2024.09.29.615696

**Authors:** Braison P. Mkiwa, Ernest W. Mauya, Justo N. Jonas, Gimbage E. Mbeyale

## Abstract

Despite the numerous negative effects of tropical forest fires in Tanzania, the causes and effects remain insufficiently documented. This study aimed to develop an integrated approach of using geospatial tools and socio-economic data to assess and map forest fire burn severity in West Usambara Mountain Forests. Three approaches including Participatory Rural Appraisal (PRA), satellite image analysis, and direct observation were used to generate information on spatial and temporal forest fire severity. Findings revealed that agricultural activities (44.5%) and charcoal production (21.1%) are the primary causes of these fires. Burn severity maps were created using the differenced Normalized Burned Ratio (dNBR) index, and the combined high and low severity areas ranged from 32.120% to 20.31%. The differenced Normalized Difference Vegetation Index (dNDVI) maps showed the combined high and low severity areas of 36.30% to 21.10% of the total study sites respectively. Post-fire NDVI time series analyses revealed a sharp decrease from 0.21 to 0.36 in all burned areas, indicating significant vegetation loss. Therefore, this study demonstrated the potential of integrating Remote Sensing and Socio-economic aspects highlighting the need for improved forest fire management through sustainable practices that balance economic and ecological considerations, offering insights that can be upscaled to other forest areas for effective management.

## Introduction

Tropical forests are recognized worldwide for their importance in the protection and preservation of various ecosystem services. For example, according to [1], tropical forests alone store a quarter of a trillion tons of carbon above and below biomass [2]. Unfortunately, they are deliberately affected by forest fire dynamics, causing approximately 30% of degradation in tropical forests caused by high temperatures and low humidity [3]. With increased forest fires in recent years, the flora and fauna in these ecosystems have been adversely affected in multiple ways through human and natural activities [4].

Shifting cultivation remains the main practice of indigenous people in many tropical parts of the world [5]. Likewise, the Convention on Biological Diversity (CBD), Aichi Target 11 emphasizes the necessity for effective and equitable management of protected areas [6], highlighting the importance of understanding local community perceptions regarding the causes and effects of forest fires since perceptions can significantly influence the success of conservation efforts [7]. Burned forests become severely damaged, causing the loss of native natural forests, introduction of invasive species that pose a significant threat to the native species, and ultimately cause massive economic loss to the people [8].

Forest fires are a major problem and protecting them is our concern [9], as they are linked directly or indirectly to intentional or unintentional human factors [8]. Meanwhile, Tanzania like many countries in sub-Saharan Africa, experiences an annual burning of about 403,400 ha of land, with approximately 12% of its land lost to fire each year between 2001 and 2007 [10-11], ranking the country fourth in SADC [12]. Miombo woodlands contribute to 75% of these fires, followed by forest plantations (20%), and forest reserves (5%) [9].

On the other hand, due to the climate changes that are currently facing the world, tropical montane forests such as Magamba Nature Forest Reserve (MNFR), Mkusu Forest Reserve (MFR), and Shagayu Forest Reserve (SFR) in the Western part of EAMs of Lushoto District in Tanzania are also facing a similar problem of frequent forest fires. It is reported that between 1997/1998, 2016, and 2021/2022 about 6,110 ha in the total area were burned in Magamba NFR [13] while, about 210 ha, and 120 ha for MFR and SFR respectively also were burned in 2021/2022 [14]. This indicates that forest fires have a significant impact on forests and can cause severe substantial environmental harm if not properly managed [9].

Despite the numerous negative effects of fire, the recent comprehensive research on the causes and effects of forest fires in tropical rainforests, particularly in the West Usambara of EAMs in Tanzania remains insufficiently documented [11,15]. Furthermore, preventive and mitigative measures for minimizing the ever-increasing threat of forest fires are inadequate to identify the long-term fire monitoring programs in Tanzania through mapping, size estimation, and distribution of forest fires [5,12]. In particular, uncontrolled fires in tropical EAMs pose a significant threat because the plants in these ecosystems are not adapted to fire events as a result promote biological invasion that sometimes are extremely flammable and prone to fire, for example, the Blacken ferns (*Pteridium aquilinum*) which immediately become dominant in burn areas after the fire [8,16].

Because most forest fires occur in remote areas, they go undetected as a result, remotely sensed satellite data is the best alternate source for forest fire studies [5]. So through RS, continuous information over large forest areas being affected by forest fires in terms of area and time was assessed using near-infrared (NIR) and short-wave infrared (SWIR) bands from pre- and post-fire satellite images [17]. Likewise, the Landsat and Sentinel-2 satellites offer advantages in forest fire detection due to their spatial and temporal resolution [18]. Remote sensed satellite data are used to create indices that indicate many aspects of the earth’s surface, such as vegetation, temperature, and humidity [5]. The Normalized Difference Vegetation Index (NDVI) is a potential index for assessing vegetation status, detecting burnt areas, and changing flora due to forest fires [19]. The Nomalize Burn Ratio (NBR) and differenced NBR (dNBR) are also included in this study since are widely used to infer fire severity from remotely sensed data [20].

The study aims, to develop an integrated approach of using geospatial tools and social-economic data to assess forest fire severity and accurately map burned areas in West Usambara Mountain Forests. Specifically to i) Map spatial distribution of forest risk zones and severity burned areas ii) Determine spatial forest fire trends over 10 years, and iii) Assess stakeholders’ perception on sources and effects of fire. The study starts by identifying and assessing Forest Fire severity and burned areas by mapping them through remote sensing technology over the last 10 years from 2013 to 2023 by defining all the possible causes and effects that arise from forest fires. Afterward, the findings will be presented to the forest departments to adopt potential preventive measures and policymakers which is useful in deciding the problem of forest fires.

## Materials and methods

### Study area

The study area covers three locations namely MNFR, MFR, and SFR (Fig.1) which are part of EAM in Usambara Western Block Mountain Forests. MNFR is found in both Lushoto and Korogwe, whereas a smaller part is located in Korogwe [21]. The district was chosen based on the prevalence of fire incidences during dry season and because the local communities are actively involved in protecting the forests against destruction by forest fires [22]. Geographically, the district is situated in the Northern part of Tanga region between 4^°^ 57’ 54’’ latitude South of equator and 38^°^ 30’ 51’’ longitude East of Greenwich [13].

**Fig 1.**
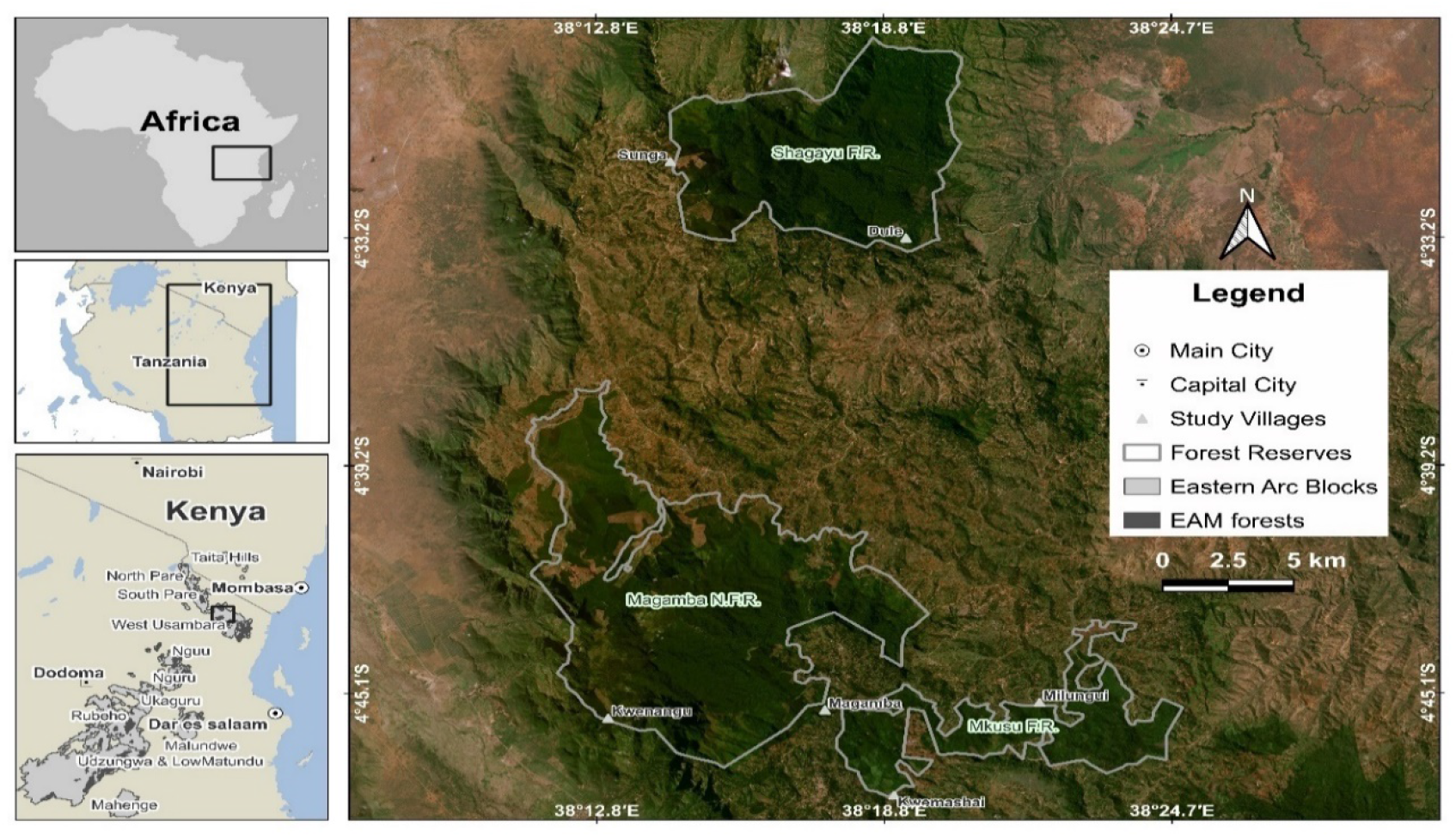
The geographical location of Magamba NFR, Mkusu FR, Shagayu FR, and study villages

The study area is located in the Tropical climatic zone and range in altitude from 1000 m to 2100 m. The District receives rainfall on a bimodal pattern, with short rains from October to December, and long rains from March to June with annual rainfall ranging from 800-2000 mm per year [21]. Temperature ranges from 15 ^0^C to 30 ^0^C annually [13]. The soils are classified as Luvic phaezem, Chromic Luvisol, Mollic Glaysol, and Rhodic Ferrasol for various activities and crops such as maize, beans, fruits, sweet and round potato, spices, and vegetables to mention a few. Table 1 summarizes the geographical location of each study forest reserve.

**Table 1.** Geographical location of MNFR, MFR, and SFR in Lushoto and Korogwe Districts.

| Name of the FR | District | Size (Ha) | Latitude | Longitude | Altitude (m) | Rainfall (mm) | Temperature (°C) | GN |
| --- | --- | --- | --- | --- | --- | --- | --- | --- |
| Magamba NFR | Lushoto & Korogwe | 9,283 | 38°15'E | 40°40'S | 2,800 | 1,200 | 22.5 | 103 |
| Mkusu FR | Lushoto | 3,674 | -4°46'E | 38°22'S | 1,600 | 900 | 26 | 114 |
| Shagayu FR | Lushoto | 7,830 | 38°16'E | 43°10'S | 1,668 | 1000 | 20 | SCH |

### Research design

The study adopted a cross-sectional and mixed research design to collect qualitative and quantitative data through satellite images and socio-economic data, as presented in the methodological flowchart in Figure 1 below. Qualitative data was collected through PRA method while quantitative data was collected using questionnaires. Key informant interviews, Focal Group Discussion (FGD), household interviews, and preference ranking (for matrix ranking) are among the PRA tools used.

### Data collection methods

#### Socio-economic data

The field survey started on December 15, 2023, and ended on February 29, 2024, in Lushoto and Korogwe Districts. Participatory Rural Appraisal tools were used in gathering social-economic data by developing semi-structured questionnaires [23]. Purposive sampling was used to select two villages from three pre-selected forest reserves based on the fire incidence and distance from the forest. Key informants including Magamba NFR staff, DFO, individuals knowledgeable about forest fires, environmental officers, village chairpersons, Village Executive Officers, and Village Natural Resource Committees (VNRC) were interviewed [23]. Additionally, questionnaires were administered to household heads during the data collection process covering general information on the local community perception on the causes and effects of fire, and how the fire regime has changed in the study areas in the last 10 years between 2013/2023. Respondents were determined based on the community who were directly involved in forest use, fire management, or handling fires. The sample size of 100 (10%) households was considered in the study, 62 were males and 38 were females from 1004 households in six study villages [24]. A checklist of questions was developed to guide the interview and discussion with the key informants. Also, household information regarding member’s age, gender, marital status, income sources, educational level, land ownership, number of livestock, and employment were obtained.

Alternatively, the procedure for obtaining the household sample size in each village involved randomly selecting 10% of household heads from the total number of households (N) [25]. The total number of households (N) in each village was divided by 10% to determine the sample size for that specific village (Table 2).

**Table 2.** The Households and sample size.

| Forest Reserve | Ward name | Village name | Total population | No of Household | Village sample size 10% |
| --- | --- | --- | --- | --- | --- |
| Shagayu FR | Mlalo | Dule M | 952 | 81 | 8 |
|  | Mtae | Sunga | 2,000 | 110 | 11 |
| Magamba NFR | Magamba | Magamba | 12,232 | 411 | 41 |
|  | Mkumbara | Kwenangu | 3,657 | 131 | 13 |
| Mkusu FR | Migambo | Milungui | 6,447 | 210 | 21 |
|  | Kwemashai | Kwemashai | 583 | 61 | 6 |
| **Overall Total** |  |  | **25,871** | **1,004** | **100** |

Respondent’s perception was elicited by obtaining their satisfaction ranking on the causes and effects of fire as an approach toward forest management. The respondents were specifically asked to rank the causes and effects of fire based on the Likert scale using five points ranging from 1 (Not known) to 5 (very well known) concerning the management aspects.

#### Remote sensing data

The first step of the method consisted of gathering, extracting, and producing the necessary data sets of satellite images for the data research area obtained by GEE. In image selection, the seasonal occurrence of fire and the absence of clouds were taken into consideration during data collection on GEE for the study areas. For this purpose, image datasets were collected before and after the fire for the period of 2013 to 2023. NBRpre, NBRpost, dNBR, NDVIpre, NDVIpost, and dNDVI indices were calculated on the images with the median statistics on the GEE cloud platform. Additionally, time series analysis was performed by calculating NDVI and NBR for each satellite image created.

### Data analysis

#### Socio-economic data

Quantitative socio-economic data collected through questionnaires were coded and processed using IBM-SPSS version 27 statistical software, while the analysis was carried out through descriptive statistics and R software to generate frequencies and percentages presented in tabular forms. Qualitative data from the PRA exercise and key informants were analyzed by using content analysis to suffice qualitative evidence. Multiple response analysis was also performed to determine responses and percentages of respondents.

#### Remote sensing data

Data was analyzed using the GEE cloud platform. Since fire alters the spectral properties of the land surface by reducing vegetation and moisture content, therefore leads to decreased reflectivity in the visible and near-infrared wavelengths, while shortwave infrared reflectivity increases [18]. Several approaches used to assess forest burn mapping using satellite-based parameters were adopted to analyze forest fires [26]. The Normalised Burn Ratio (NBR) spectral index developed by the US Geological Survey (USGS) is a widely used approach for this purpose [17]. The NBR compares the near-infrared (NIR) and short-wave infrared (SWIR) reflectance values, with healthy vegetation showing high NIR reflectance and low SWIR reflectance [20]. This index is effective in distinguishing between areas affected by fire and areas with healthy vegetation based on their spectral characteristics and ranges from -1 to +1 [17].

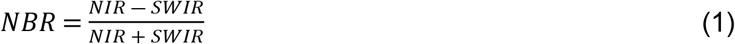

In contrast to bare land and recently burned areas, high NBR values indicate healthy vegetation. Non-burned areas are normally attributed to values close to zero [26]. The difference between the pre-fire and post-fire NBR is used to calculate the delta NBR (dNBR), which can be used to estimate burn severity. The dNBR ranges from -2 to +2 with high positive values representing severely burned areas [17]. The higher dNBR values indicate more severe damage, while negative dNBR values indicate regrowth following a fire [20]. It is therefore the appropriate index for discriminating between burned and unburned areas, which contains information in the NIR and SWIR spectrum regions.

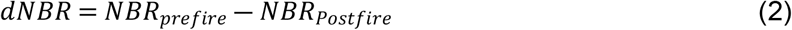

The dNBR index of the study area was calculated for two periods corresponding to the forest fires in 2013 and 2023. Therefore, four Landsat datasets (2014, 2017, 2020, and 2023) were used to get NBR and NDVI values (pre- and post-fires).

The NDVI can predict vegetation and biomass change pre-fire and post-fire, its values range between -1 and 1 [27]. Vegetated areas take positive values, while the negative values correspond to bare soils. High NDVI values represent dense green areas such as forests and cultivated areas [19]. NDVI equation was used to calculate the vegetation index to estimate the spatial and ecological biomass changes before and after the fire as well as an equation to obtain pre-fire and post-fire differences [27].

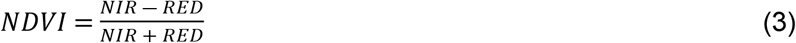

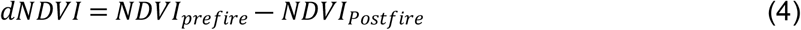

In this study, the dNDVI index was calculated by subtracting the pre- and post-fire NDVI values (Eqn 4), and its values were reclassified into five classes (Fig. 7).

## Results

### Socio-economic data

#### Socio-economic characteristics of the respondents

The average age of the respondents is between 31 and 40 years, with an annual household income of 2,422,216 TZA. About two-thirds (70.50%) of the participants were male and the remaining (29.50%) were female (Table 3). On the other hand, about 51.15% of the respondents completed primary school, 31.82% completed secondary school level, 17.04% completed higher education level, and none for informal education. Agriculture activities made up a vast majority (78.40%), followed by those working in beekeeping activities (8%), and the list one being mason (1.10%) as summarized in Figure 3.

**Table 3.**
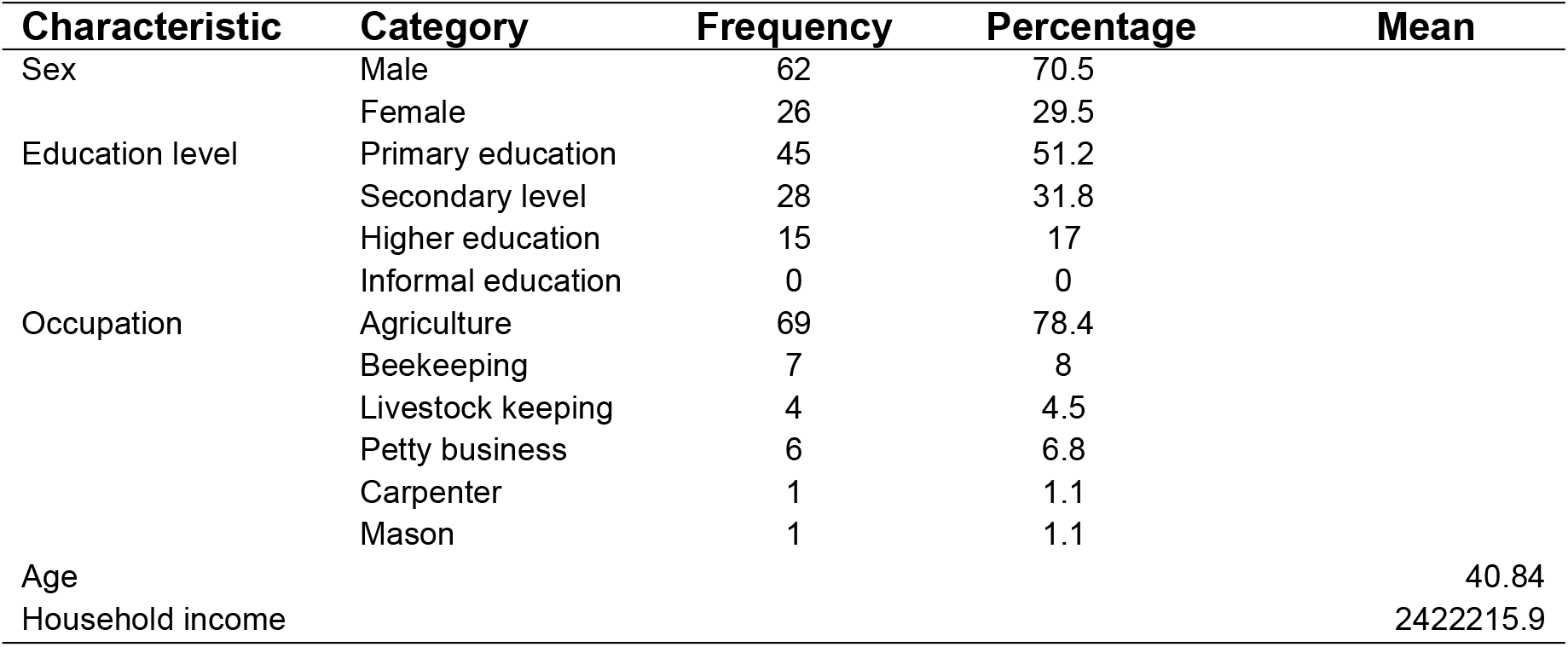
Socio-demographic Characteristics.

**Fig 2.**
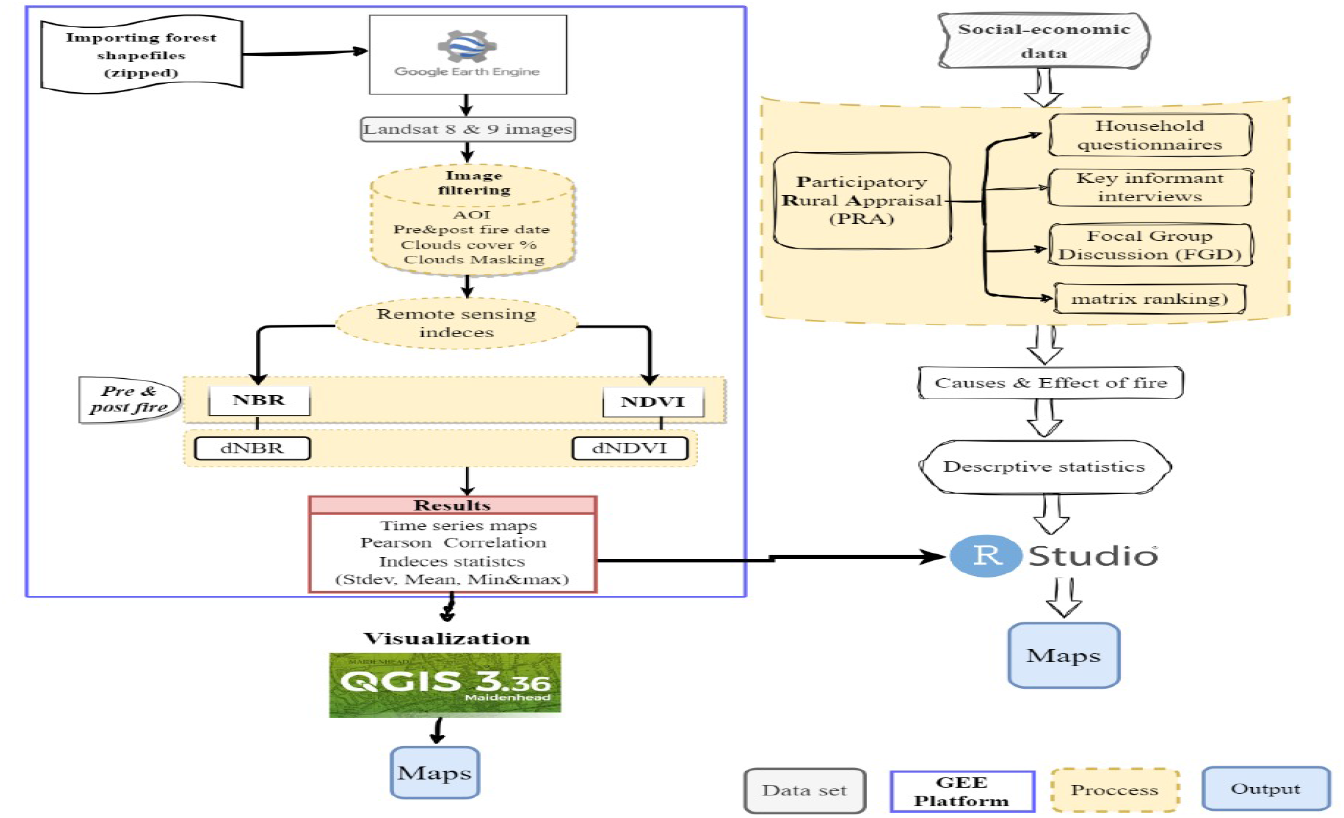
Methodological Flowchart Depicting the Link of Geospatial and Social-economic Data to Analyse Forest Fire Burn Severity in the West Usambara Mountain Forests

**Fig 3.**
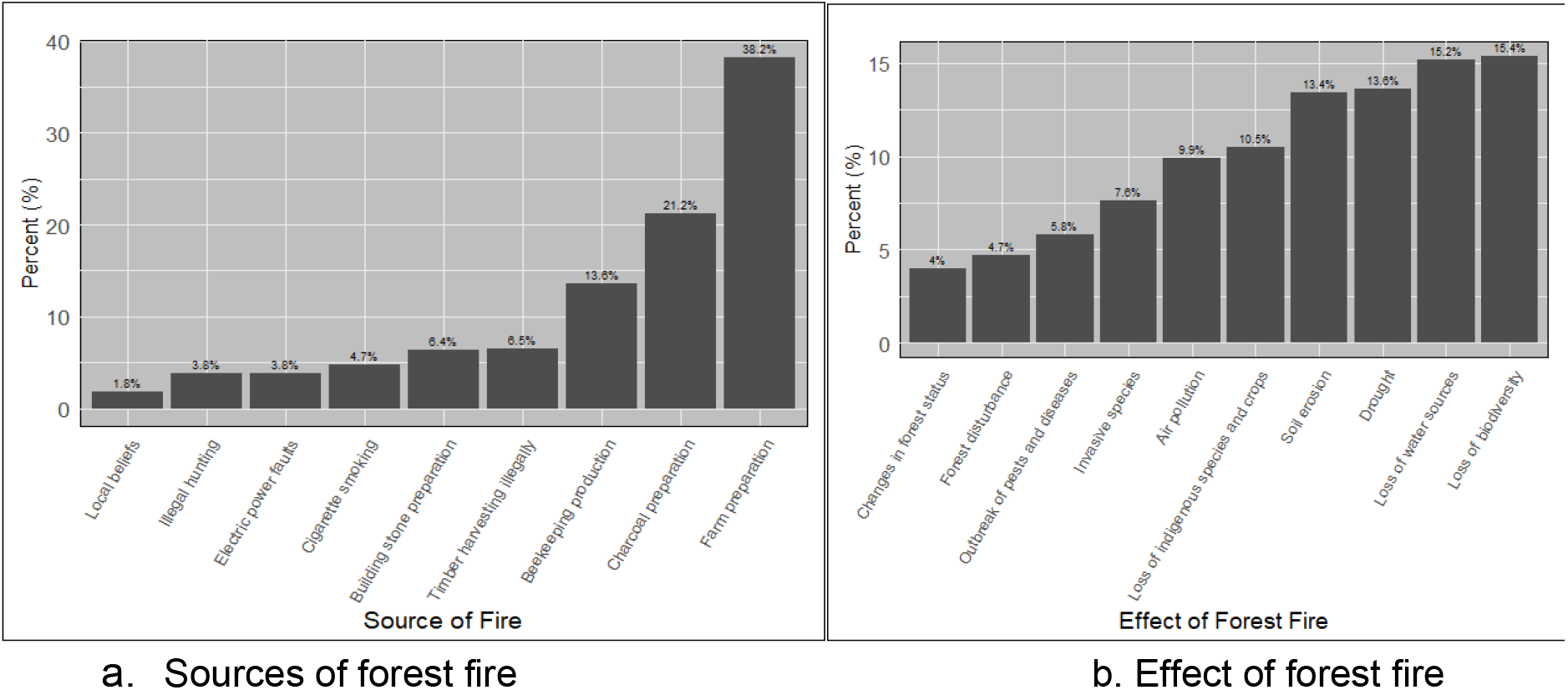
Perception of local respondents on the source (a) and effects (b) of forest in the study area

#### General Perception of local respondents on the source and effects of forest fires in the study area

The sources of bushfires in Lushoto District are illustrated in Figure 3, with farm preparation identified as the leading cause at 38.2%, while local beliefs accounted for only 1.8%. Respondents ranked various effects of forest fires, with the loss of biodiversity being the most significant concern at 15.4%, followed by changes in forest status at 4% as summarized in Figure 3. Additionally, respondents ranked their perception and knowledge of forest fire from six topics describing how they understood and perceived the complexity of managing fire where (60%) of the respondents know the presented topics as illustrated in Figure 4.

**Fig 4.**
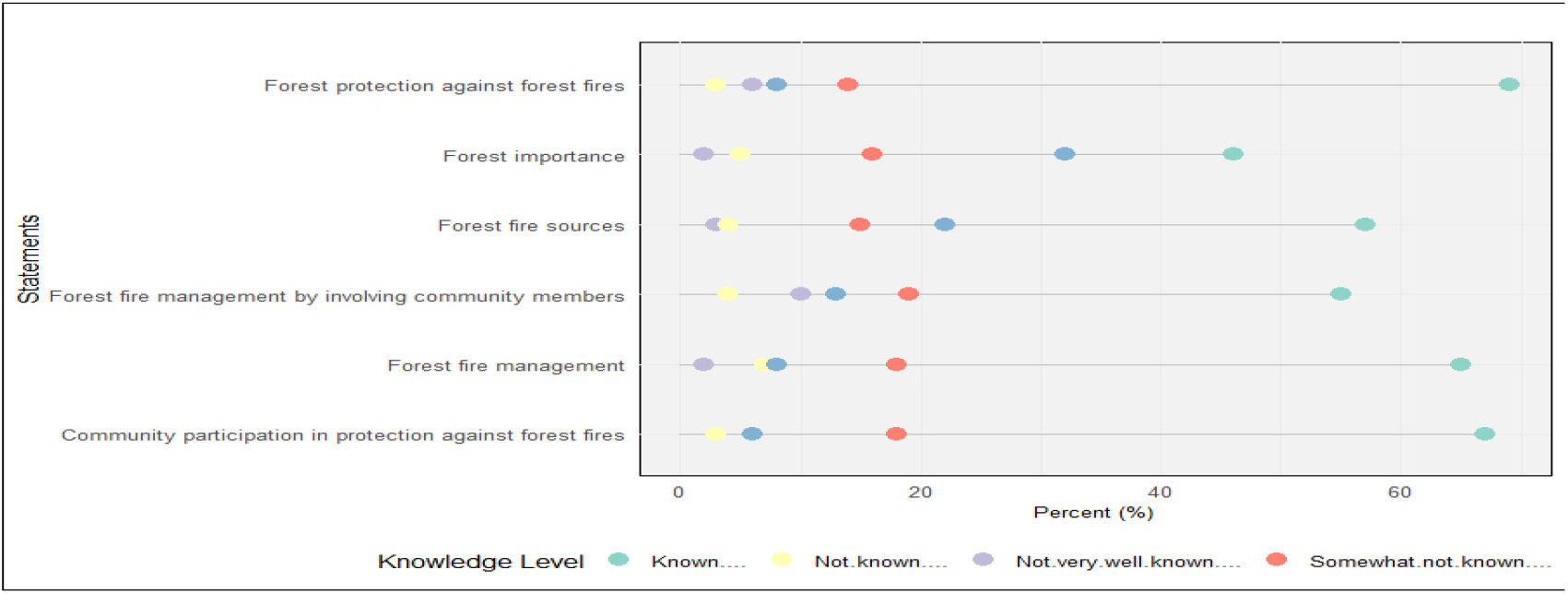
Likert scale rating regarding perception and knowledge topics on forest fire from respondents.

#### Remote sensing data

##### Mapping burned areas

Detecting forest fire areas using Landsat-8 and 9 satellite imagery and its vegetation indices has become a potential method for estimating forest fires [20]. The NBR index derived from the sentinel-2 satellite image was used to characterize burned areas and assess forest fire severity. At the same time, the NDVI variables confirm vegetation status within the forest over time. The NBR mean values ranged from 0.20 to 0.32 (Fig. 6) and were categorized into five classes (low severity, high severity, unchanged, low regrowth, and high regrowth) (Fig. 7). The spatial analysis from the map reveals that from 2013 and 2023 fire period the study area was dominated by low (20.06%) and high (32.19%) burn severity. For this purpose, dNBR values were calculated from NBR_pre-fire_ and NBR_post-fire_ (Eq.2) using Landsat-8 and 9 images produced in the study area of 30 m spatial resolution from the GEE cloud platform.

**Fig 5.**
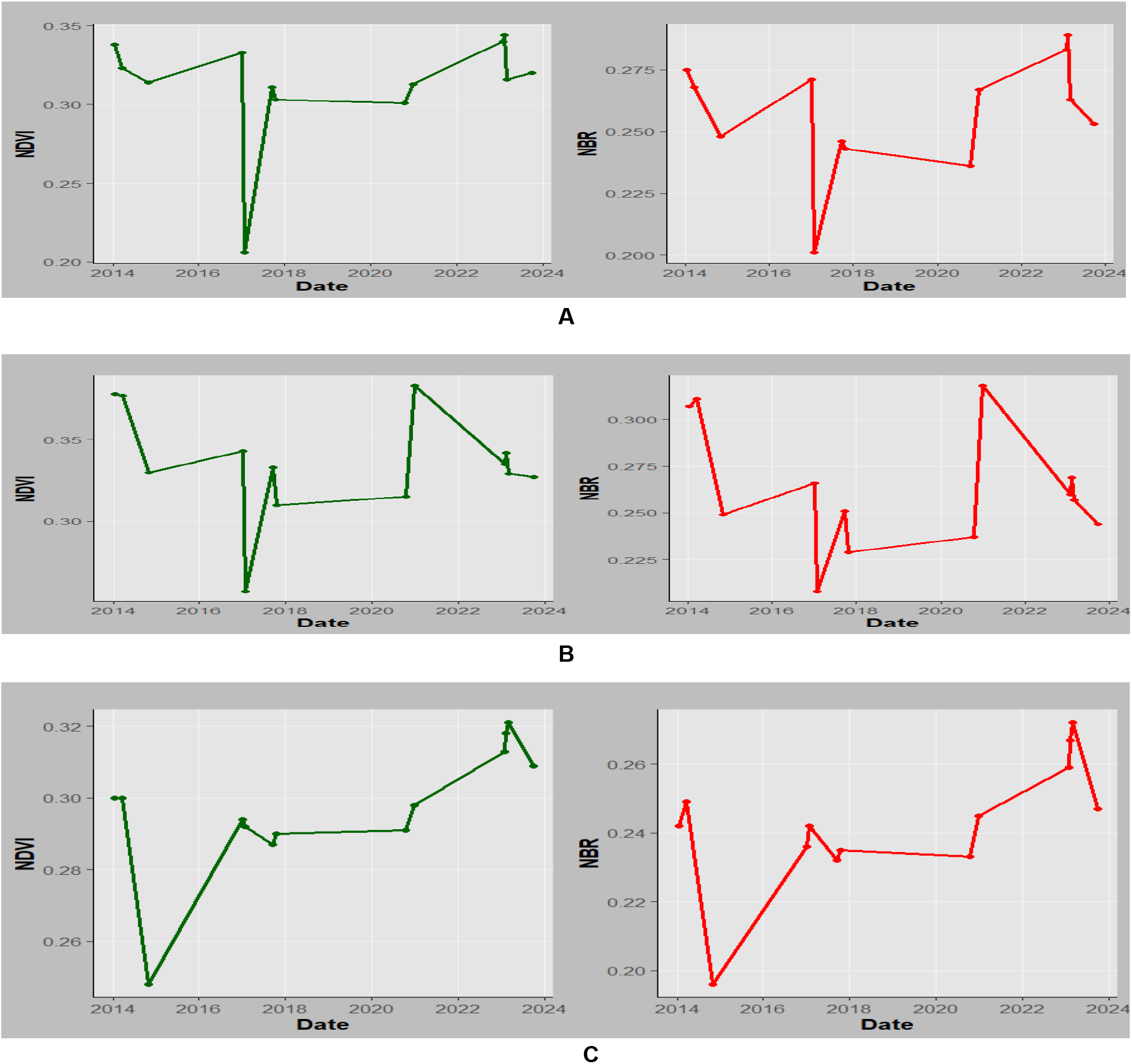
NBR and NDVI time series for (a) Mkusu FR (b) Shagayu FR, and (c) Magamba NFR

**Fig 6.**
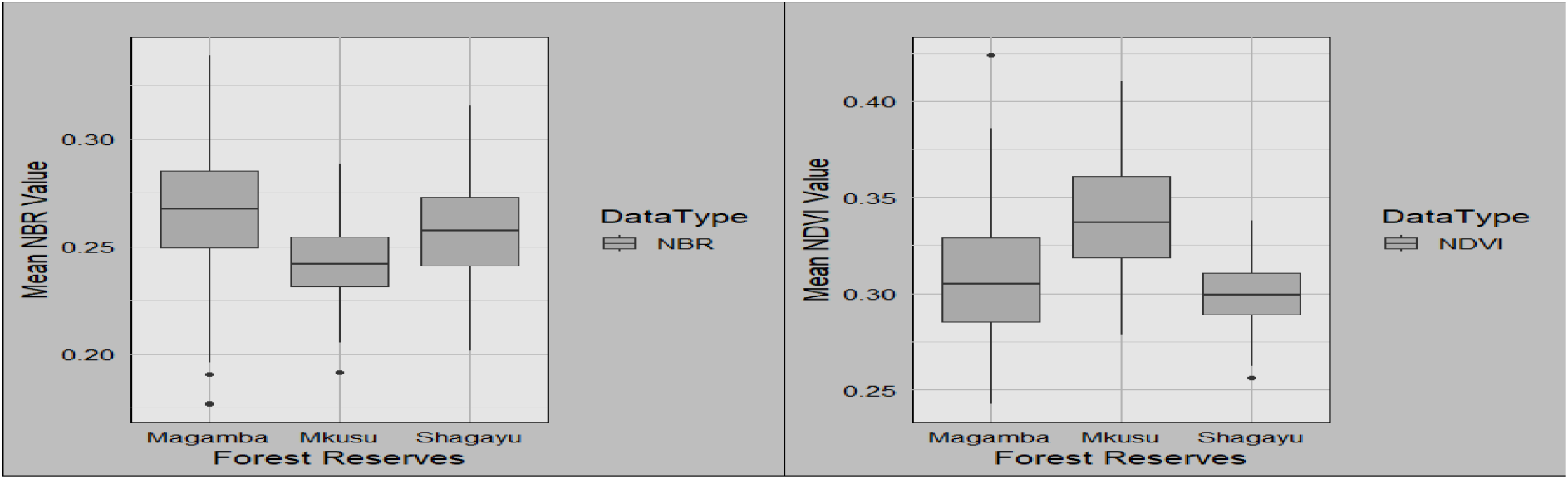
The NBR and NDVI mean values for Magamba NFR, Mkusu FR, and Shagayu FR.

**Fig 7.**
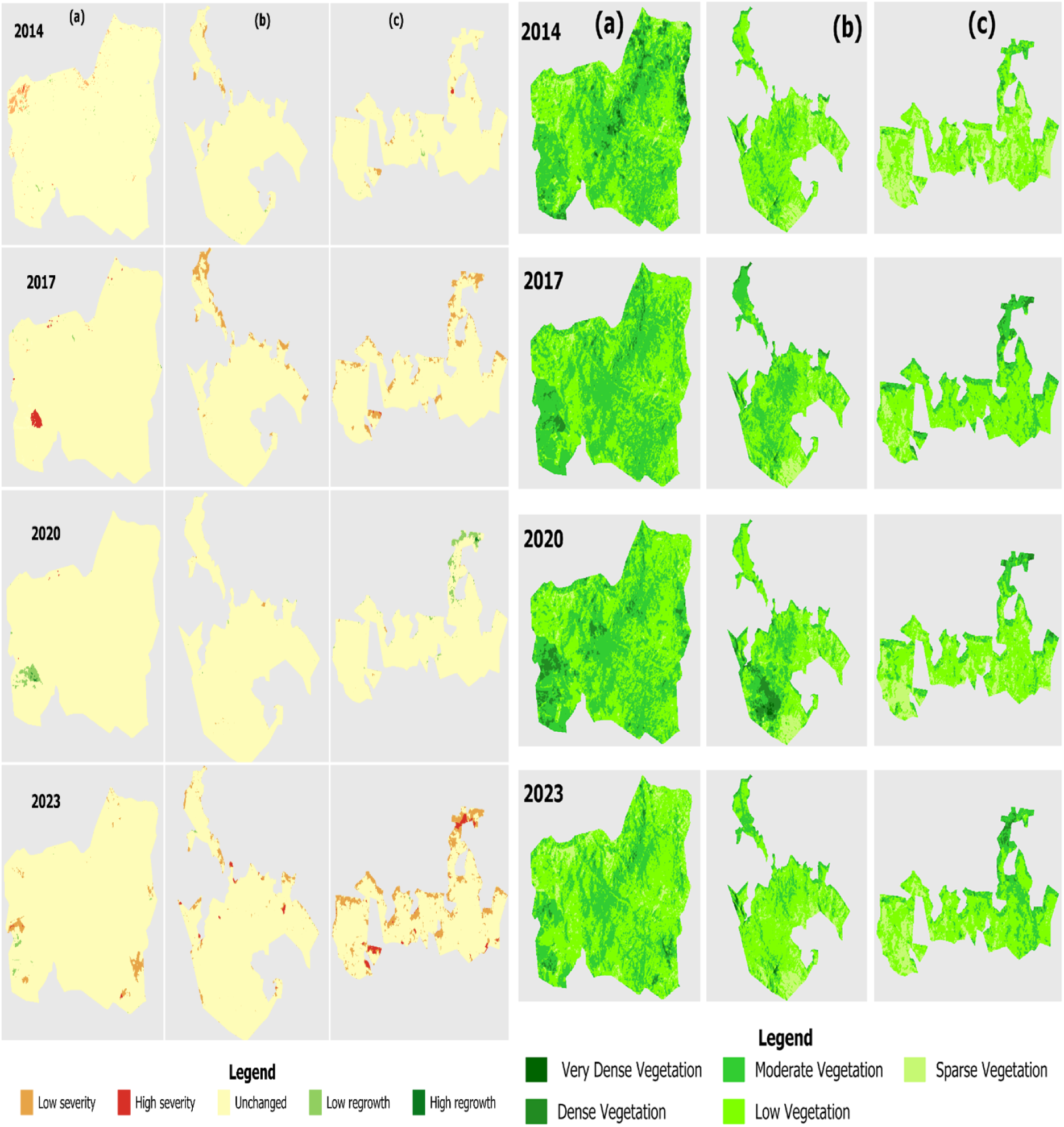
Spatial distribution of burn severity maps created according to NBR-left and NDVI-right (a) Shagayu FR, (b) Magamba NFR, and (c) Mkusu FR

NDVI maps are classified as very dense vegetation, dense vegetation, moderate vegetation, low vegetation, and sparse vegetation (Fig.7). Additionally, the non-burned areas from the 2013 and 2023 fires represented 11.47%, 20.4%, and 30.09% of the study area in Magamba NFR, Mkusu FR, and Shagayu FR respectively (Fig. 9).

**Fig 8.**
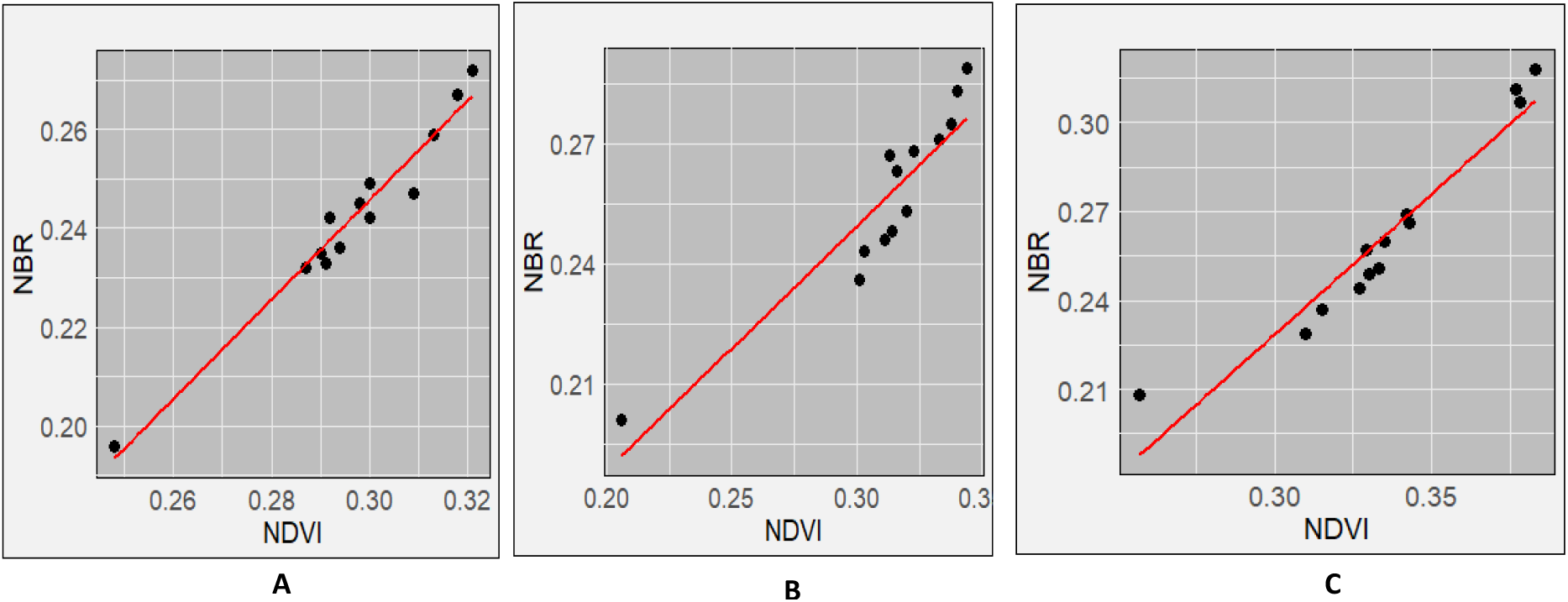
Correlation analysis between NBR and NDVI for (a) Shagayu FR, (b) Magamba NFR, and (c) Mkusu FR (c).

**Fig 9:**
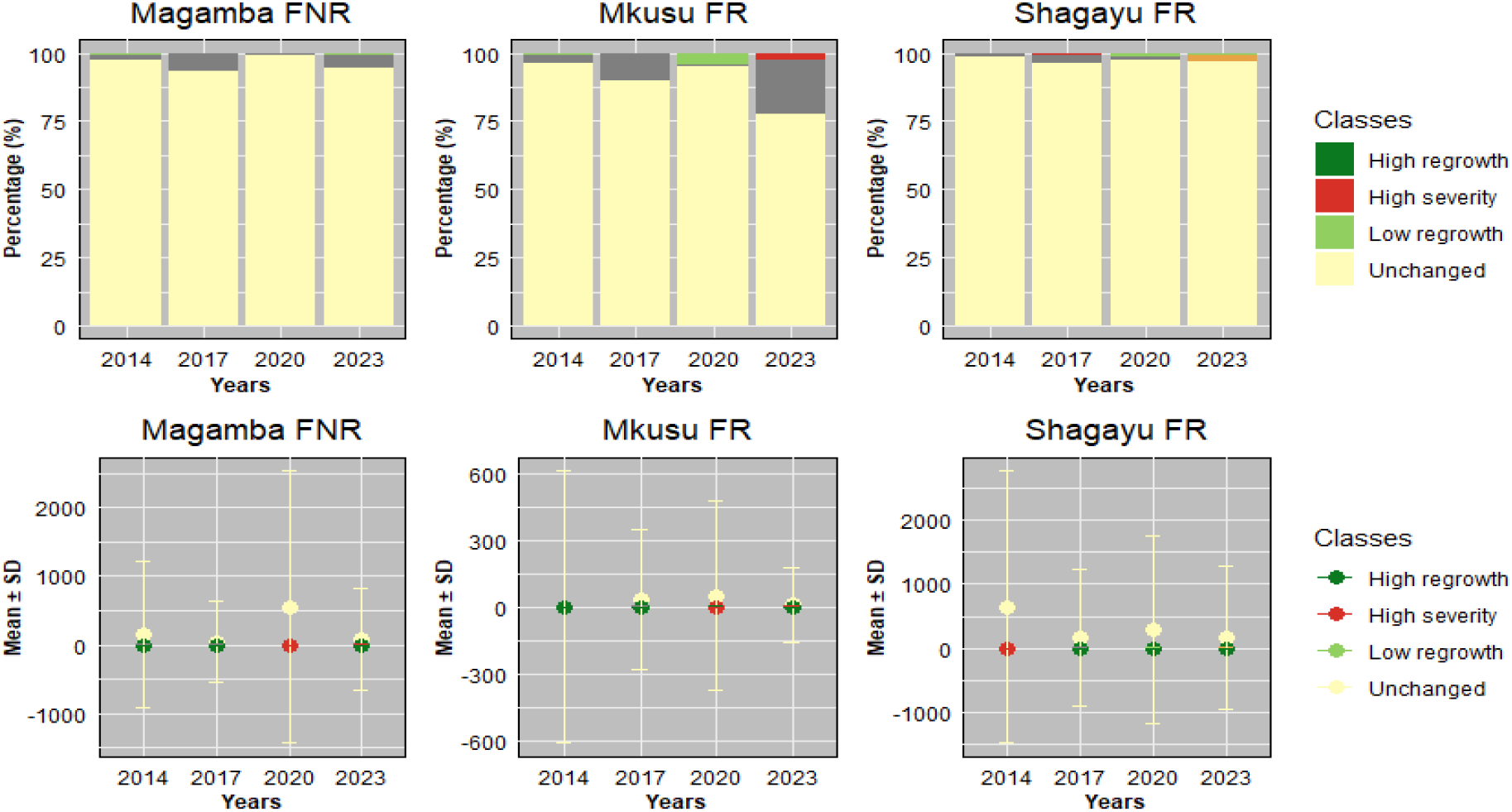
Area changes over time for Magamba NFR, Mkusu FR, and Shagayu FR.

The total burned area was 3,296.96 ha, which constitutes 15.86% for both Magamba NFR, Mkusu FR, and Shagayu FR (Fig. 9). Areas with low and high regrowth and unchanged account for 84.14% of all total areas. In Magamba NFR, the fire affected 1322.80 ha, 84.14% of all total regions. In Magamba NFR, the fire affected 1322.80 ha, which is 14.25% of the total area, with low and high-severity fires impacting 1272.05 ha and 50.75 ha respectively. Mkusu FR experienced a total burned area of 1341.23 ha, which is 36.50% of the total area with high and low-severity fires affecting 108.88 ha and 1232.35 ha respectively. Shagayu FR had a burned area of 632.93 ha, which is 8.1% of the total area with 575.04 ha and 57.89 ha for low and high-severity fires respectively. Mkusu FR was identified as the most affected reserve, while Shagayu FR was the least affected.

The correlation between the NBR and NDVI was calculated in each Forest Reserve. Their values for each point were then computed using Microsoft Excel to plot the scattered charts shown in Figure 8. Table 4 shows the correlation coefficients R (0.92, 0.96. and 0.98) and their respective R^2^ values for Magamba NFR, Mkusu FR, and Shagayu FR respectively. These statistics values indicate strong positive linear correlations between NBR and NDVI indices, which are compared to assess fire severity and vegetation condition derived from RS data and field observations. Shagayu FR shows the strongest relationship, followed by Mkusu FR, and then Magamba NFR.

**Table 4.** Correlation coefficient (R) and coefficient of determination (R^2^) values in the study area.

| Forest name | Correlation Coefficient (R) | Coefficient of Determination (R<sup>2</sup>) |
| --- | --- | --- |
| Shagayu | 0.98 | 0.96 |
| Magamba | 0.92 | 0.84 |
| Mkusu | 0.96 | 0.92 |

Meanwhile, when assessing the fire severity and vegetation health between NBR and NDVI indices in Figure 8, it is evident that as NBR values increase, NDVI values also tend to rise in both study areas. It suggests that as areas recover from fire damage (indicated by higher NBR values), the health of vegetation (as reflected by NDVI) also improves. Therefore, the strong relations between NBR and NDVI suggest the potential use of these spectral indices for burn severity estimation and monitoring post-fire recovery. The mean NBR values indicate the severity of the burn while the mean NDVI values provide insight into the health and recovery of vegetation. For example, in Figure 6 the NBR mean values of Magamba NFR, Mkusu FR, and Shagayu FR were 0.26, 0.26, and 0.24 respectively.

Likewise, NDVI and NBR mean values in Magamba NFR, Mkusu FR, and Shagayu FR were 0.31, 0.34, and 0.30 respectively (Figure 6). The high NDVI mean values indicate dense, healthy vegetation while the low NDVI mean value reflects sparse or unhealthy vegetation.

##### NBR and NDVI Time series analysis

The time series assessment of both NBR and NDVI performed in the GEE platform often shows similar trends with little deviation with some points in response to fire events. For instance, both indices typically exhibit a decline immediately after the fire (2015, 2017, 2018, and 2024 for both MFR and MNFR), followed by gradual recovery as vegetation begins to regrow between 2016, 2019-2023 (Fig. 5).

## Discussion

The study aimed to develop an integrated approach using geospatial tools and socio-economic data to assess and map forest fire burn severity in the West Usambara Mountain Forests. RS and socio-economic data reveal that agriculture and charcoal production are the primary causes of forest fire burn severity.

The findings of this study indicate that the average age of the respondents between 31 and 40, reveals a significant presence of young, financially capable individuals. At the same time, those with 60 and above are less active economically [23]. Meanwhile, African traditional gender roles highlight distinct social roles assigned based on gender, where men typically head households [28]. Additionally, higher education levels are associated with increased access to technical information on fire management, enabling individuals to adopt innovative and sustainable practices [29], as it is hypothesized that as education levels rise, more respondents will embrace sustainable fire use practices advocated by CBFiM [30]. Other studies have also found that older people perceive conservation positively as a gift for future generations [31].

The widespread practice of zero grazing in the study area suggests community awareness of land management practices [23]. However, using fire for farming and charcoal production contributes significantly to forest fires [9,15]. Forest fires adversely impact biodiversity, which is vital to the community, affecting traditional knowledge, economic stability, food security, and cultural practices, as reported by key informants. The perception of more high-intensity fires in the study area over the past decade is supported by the literature on the case of increased burned areas in northwest Portugal [32]. These fires also influence ecosystem services, recreational values, and educational opportunities [15]. In particular, agriculture is a significant source of income for villagers in the study area, with common crops including maize, beans, potatoes, and vegetables. The prevalent practice of slash-and-burn agriculture, especially in areas experiencing population growth, significantly contributes to the occurrence of forest fires in tropical fires [11,15,33].

Furthermore, charcoal production and cultural beliefs about fire use can contribute to forest fires, although these are less prevalent in the study area. For example, in Sunga village, one informant stated that *‘‘Sambaa tribe believes that if one starts a fire and it ends up getting and spreading to a large extent, signals a long life’’*. Local respondents recognize the importance of fire for various activities but also acknowledge the devastating consequences of unmanaged fires, such as loss of biodiversity and forest degradation [15]. Despite community awareness and education efforts, high-fire incidents continue to occur in national forest reserves in Tanzania, [34]. According to the key informants, the availability of significant tangible benefits such as economic incentives from eco-tourism, water for irrigation, and domestic use are the reasons for active willingness to be involved in forest fire protection similar to these scholars [35-36]. Lack of or insufficient tangible benefits from national forests can lead to disappointment among local people [37].

Findings from the Remote Sensing component of this study confirm that reliable burn severity estimates can be derived from combined indices of NBR and NDVI for mapping burn severity and vegetation status. Through NDVI and NBR indices, we tested and compared for burn severity mapping to justify the results (Fig.7), as [38] did in their study research. The NBR index records the effect of fire with enhanced spectral contrast, making it easier to estimate burned areas, assess burn severity, and monitor vegetation recovery after a fire [17]. This study demonstrated high accuracy in detecting fire-affected areas using the NBR and NDVI indices derived from Sentinel-2 images, as supported by multiple research findings [17,26,27].

Similarly, the NDVI was used to detect the extent of vegetation status and destruction of existing vegetation before and after the fire as supported by [39]. When comparing NBR and NDVI results, the NDVI results analysis observed that areas burned transformed into open spaces or bare areas within the forest after the fire as supported by [39]. According to [40], the healthier and lusher the vegetation, the higher the corresponding NDVI value. [26] used the dNBR index derived from Sentinel-2 images to detect forest fires similar to the approach taken in this study. While dNBR is effective for detecting burned areas, it is sensitive to water, which can lead to misclassification of high-severity pixels; however, water masking was applied in the study area to address this issue [41].

Consequently, despite the numerous studies that show strong correlations between field-based reference observation of burn severity and both post-fire and differenced spectral indices [27,42,43], the sensitivity of these methods varies across different vegetation types [39], so they need to be field-validated to ensure that reliable information is generated. The strong correlation enhances the reliability of using these indices for effective fire management and assessment with R^2^ values reflecting the effectiveness of RS data in accurately estimating fire effects [20]. The strong correlation between NBR and NDVI values typically indicates a strong positive correlation as vegetation health improves (higher NDVI), and the area is likely to recover from fire damage (higher NBR) [44]. On the other hand, the mean NBR values indicate the severity of the burn while the mean NDVI values provide insight into the health and recovery of vegetation [20]. The high NBR mean values indicate healthy or unburned vegetation while the low NBR mean value indicates that the area has experienced significant fire damage [27]. Also, the high NDVI mean values indicate dense, healthy vegetation while the low NDVI mean value reflects sparse or unhealthy vegetation.

Similarly, findings further show that a sharp decrease in NBR and NDVI in 2017 for MFR and SFR indicates forest burning during that period. This correlation reinforces the reliability of using both indices for monitoring fire effects and assessing vegetation status [27]. The high correlations suggest that changes in one index can help to infer changes in the other index, making them valuable tools for forest fire management and ecological studies [41].

Additionally, NDVI time series analysis is used to track changes in vegetation, particularly in areas that have been rehabilitated following wildfires [27]. It is commonly used to identify vegetation density changes before and after fire. The observed decrease in NDVI in 2017 in this study aligns with [19] who found the same scenario in Arouca. [45] reported that post-fire NDVI values decreased to 0.36 in areas with high burn severity while for this case, a sharp decrease up to 0.21 in Mkusu FR was observed. [46] examined the dynamic behavior of pre and post-fire vegetation in a burned forest area through NDVI time series analysis and they found a sharp decrease in post-fire NDVI values in Northern Italia. Also, [47] found similar results for Madeira Island when detecting forest fire severity and vegetation health over time using NBR and NDVI indices also similar to this study. In this study, the areas where the fire occurred are highly dominated by Blacken ferns (*Pteridium aquilinum*) which immediately become dominant in burn areas after fire [16]. After the fire, NDVI values were between 0.21-0.36, indicating a decrease in the dense vegetation. Therefore, low NBR and NDVI mean values show substantial burn and loss of vegetation health and density in some periods as the study aligns with [27].

## Conclusion

The present study integrates geospatial tools and socio-economic data to identify forest fire burn severity zones, sources of fire, and their effects using Sentinel-2 satellite data and the Google Earth Engine (GEE) cloud platform to create thematic layers of burn severity and ultimately produce a comprehensive synthetic burn severity maps. The use of the GEE platform in this study significantly enhances geospatial data processing capabilities, advancing research on forest fire prediction. GEE provides a fast, reliable, and effective means of producing burn severity maps, offering substantial advantages for monitoring pre and post-fire conditions, particularly in large and mountainous forest areas. Remote Sensing findings indicate that NBR and NDVI indices effectively estimate burn severity and give clear vegetation status. On the other hand, socio-economic data indicates that incorporating stakeholder perceptions into management decisions enhances societal acceptability and effectiveness, with the community identifying shifting cultivation and charcoal production as the primary causes of forest fires. Therefore, the study provides a basis for future fire management and prevention efforts, emphasizing the need for awareness regarding the sources and effects on livelihood and the effective control mechanisms. They also highlight the critical importance of burn severity mapping for informed decision-making and strategic planning in both high and low-risk zones of the west Usambara Forest in Lushoto, Tanzania.

The study recommends enhancing community participation in forest conservation by increasing local knowledge and awareness of Community-Based Fire Management (CBFiM) and Integrated Fire Management (IFM) approaches. This involvement fosters a sense of ownership and shared responsibility, addressing encroachment and managerial challenges. Future research should explore the long-term impacts of forest fires on biodiversity and ecosystems in the Western Usambara Mountain forests.

## Acknowledgments

The authors thank the Tanzania Forest Services Agency (TFS) and Tanzania Forest Fund (TFF) for financial support for field work and manuscript development.

## Author contributions

**Conceptualization:** Braison Paul Mkiwa, Ernest Mauya, Gimbage E. Mbeyale.

**Data curation:** Braison Paul Mkiwa,

**Methodology:** Braison Paul Mkiwa, Ernest Mauya, Gimbage E. Mbeyale.

**Formal analysis:** Braison Paul Mkiwa, Justo N. Jonas.

**Writing – original draft:** Braison Paul Mkiwa,

**Writing – review and editing:** Braison Paul Mkiwa, Ernest Mauya, Gimbage E. Mbeyale.

**Visualization:** Braison Paul Mkiwa,

**Supervision:** Ernest Mauya, Gimbage E. Mbeyale.

